# Integrative proteomics and metabolomics map reveals key sectors of defense metabolism in ginger against *Pythium myriotylum*

**DOI:** 10.1101/2025.09.09.675089

**Authors:** Febina Fernandez, Lini Varghese, Vinitha Meenakshy Ramaswami, George Thomas, Anish Kundu

## Abstract

Ginger (*Zingiber officinale*) cultivation is severely threatened by rhizome rot caused by *Pythium myriotylum*. Hitherto, detail molecular machineries and metabolic pathways of defense against *P. myriotylum* in edible ginger are elusive. To elucidate the defense mechanisms, we employed integrative quantitative proteomics (TMT labelling) and high resolution untargeted metabolomics to investigate temporal defense responses of ginger during infection. Proteomics analysis revealed enrichment of cellular detoxification and secondary metabolism pathways indicating co-activation of oxidative stress management and its mediated defense metabolite biosynthesis. Key phenylpropanoid enzymes (PAL, C4H, CAD) and LOX, a regulator of jasmonic acid (JA) biosynthesis, were significantly up-regulated, correlating JA signaling with secondary metabolic defenses. Metabolomics profiling confirmed accumulation of phenylpropanoids, flavonoids, and terpenoids, including known antimicrobial metabolites such as gingerol, shogaol, curcumin, and gingerdione, alongside expression of their key regulatory enzyme was also induced. Notably, machine learning identified the monoterpenoid TGA as the top defense-associated metabolite. Functional assays demonstrated strong anti-*P. myriotylum* activity of TGA, which inhibited mycelial growth and suppressed expression of key pathogenicity-related genes (*NIP1*, *GHs*, cellulase). These findings highlight coordinated activation of detoxification and secondary metabolic pathways in ginger defense and establish TGA as a promising bio-control agent against *P. myriotylum*.

## Introduction

Soft rot, found in various crops is generally caused by devastating oomycetes *Pythium* species, in particular, *Pythium myriotylum* Drechsler (Kumar et al., 2008; Le et al., 2014) which is a soil borne and necrotrophic pathogen in nature, belongs to the order pythiales. This pathogen results in huge losses such as up to 90% crop death as pre and/or post-emergence damping-off in cultivation of several crops including tobacco, tomato, chilli pepper with a special threat to one of the most economically important spice crops, ginger in warm and moist weather (Hyder et al., 2021; Yoshila et al., 2023). It is prevalent in almost all ginger cultivating areas of the world and was reported from Australia and Fiji (Stirling et al., 2009), Japan (Ichitani and Goto, 1982), Korea (Kim et al., 1996) as well as Nepal (Nepali et al., 2000), and was initially reported in 1907 from Surat, Gujrat, India (Butler, 1907) which causes soft rot disease in the roots, stems, and rhizomes of wild as well as cultivated plants (Martin and Loper, 1999). Development of ginger varieties immune to soft rot through conventional sexual hybridization methods is not amenable due to their mandate asexuality (Hooker, 1894; Dake, 1995). Moreover, due to an exceptionally narrow genetic background (Varghese et al., 2018) clonal selection is likewise not feasible as all varieties are equally prone to soft rot. Considering their incredible ability to thrive in soil, *Pythium* species are mostly uncontrollable by chemical and cultural means (Le et al., 2014). Since, the use of synthetic fungicides is ineffective against soft rot (Martin and Loper, 1999; Le et al., 2014) and the host defence mechanism against *P. myriotylum* to render genetic resistance being unexplored (Jones and Dangl, 2006), it is important to investigate the defense mechanisms involved during plant-pathogen interaction and host’s machinery to initiate defense metabolism. Being an ideal host for *Pythium* species, ginger can be a model for elucidating defense modules including early and late phase of defense responses. In a wild ginger, *Zingiber zerumbet*, cell wall fortification, lignin biosynthesis, and salicylic acid-jasmonic acid hormonal crosstalk associated transcription factors are found to be involved along with phenylpropanoid pathway and salicylic acid mediated defence against *P. myriotylum* was recently demonstrated (Nath et al., 2018; Augustine et al., 2024), however, due to limitations of the bio resources and functional mutants molecular machineries and metabolic pathways of edible ginger, *Zingiber officinale*, involved in defence against *P. myriotylum* is poorly understood yet.

The ‘integrative multi-omics’ is an outstanding approach in uncovering plant-microbe interactions specifically plant-associated defence strategies and contributes to improved crop productivity as evidenced by recent successful research (Kimotho and Maina, 2024; Kumar et al., 2023; Liu et al., 2021; Hu et al., 2018). A comprehensive knowledge on plant-pathogen interactions can be attainable via a multi-omics approach, which may assist us to design predictive models for potential plant responses to such stress (Crandall et al., 2020). Mass spectrometry based proteomics can be extensively used to study plant-pathogen interactions to understand dynamics of proteins, post translational modifications, and biological pathways (Quirino et al., 2010; Elmore et al., 2021; Jain et al., 2021). On the other hand, with recent advancements, metabolomics has been developed as an excellent tool for elucidation of host-microbiome interactions and applied plant-microbiome studies (Hu et al., 2018; Geier et al., 2020; Wolinska et al., 2021; Getzke et al., 2023). In the current investigation, we have applied a multi-omics framework by integrating quantitative proteomics and untargeted semi-quantitative metabolomics in ginger- *Pythium* pathosystem, for mapping the core defense modules of the host’s defense during pathogen attack. This high throughput data driven ‘multi-omics’ approach also discovered major metabolic pathways, associated proteins and specialized metabolites crucial for immune-metabolism in edible ginger. Finally, by applying two machine learning models, we found out *trans*-geranic acid (TGA) as a crucial defense metabolites with strong bioactivity against pathogenicity of *P. myriotylum*. Taken together, by integrating information across multiple omics layers, our study delivered an in-depth grasp of the metabolic mechanisms behind ginger’s defense against this devastating pathogen.

## Materials and methods

### Plant material and inoculation of pathogen

*Zingiber officinale* Rosc., edible ginger variety called Varada which is susceptible to *Pythium myriotylum* causing soft rot were used for the study. The plants were raised in an insect-protected net-house at natural field conditions while they were growing in earthen pots of an appropriate size in a mixture of red-earth: river-sand: leaf compost (1:1:1). The harvested rhizome parts from the mature plants were allowed to germinate to give new plants. For the purpose of pathogen inoculation, four month old plants with a similar growth and leaf stage were chosen. The plants were infected with 48-hour-old culture mycelial (10 mm diameter) of an aggressive isolate of *Pythium myriotylum* Drechsler (RGCB N14) in the collar region, which is the junction of the aerial stem and the rhizome part, as previously described (Geetha et al., 2019). The collar region is the general entry point of this soil borne pathogen. Three to four biological replicates each for both the control and the treatment conditions were kept for the experiment. With periodic sub-culturing and virulence testing, the pathogen was grown in the potato dextrose agar, or PDA medium at optimum room temperature (25 °C) in dark condition. As controls, plants that had been mock-inoculated with plain PDA discs were utilized. The collar tissue samples had been collected at three different time intervals, notably 0 hours post inoculation (control), 12 hours post inoculation (hpi), as well as 24 hours post inoculation (hpi) with *P. myriotylum*. These two time points have been chosen as early and late phase upon infection based on pathogen colonization pattern. An inch-long collar area has been collected by cutting with sterile scalpel, flash-frozen in liquid nitrogen, and kept at -80°C until required.

#### Histopathology

Lactophenol-trypan blue staining of transverse and cross sections of the collar region was used to confirm the growth of *P. myriotylum* in pathogen-inoculated *Z. officinale* plants at 0, 12, and 24 hpi and to determine the pattern of colonization of pathogen inside the host tissue (Narayanasamy & Narayanasamy, 2011). Nikon Eclipse Ni-E (Nikon, Tokyo, Japan) microscope was used for observing and taking images of the trypan blue stained sections at 20 and 40X magnification under bright field illumination.

#### Preparation of samples for proteomics

Collected ginger tissues were powdered using a pulveriser previously kept in liquid nitrogen. Around 2 mg of this powdered tissue material was precipitated in 1 ml chilled 10% TCA/Acetone and centrifuged at 14000 rpm for 10 minutes 4°C. The supernatant is discarded and the pellets were resuspended in 1ml of chilled 2% 2-Mercaptoethanol and centrifuged for 10 minutes at 14000 rpm at 4°C. The pellet is again resuspended with 1ml of ice-cold acetone followed by centrifugation at 14000 rpm at 4°C for 10 minutes. The final pellet is allowed to dry completely and is stored at ^-^80°C. The pellet was precipitated in SDS for protein estimation and further processing. Protein amount was estimated using BCA (Bicinchoninic Acid) protein assay. From all samples 400 μg of protein were reduced, alkylated and precipitated with 5 mM DTT (Dithiothreitol), 20mM IAA (Iodoacetamide) and chilled acetone. The precipitated samples were resuspended in 50 mM TEABC (Tri Ethyl Ammonium bicarbonate) and subjected to Trypsin digestion (1:20 Promega) for 12-16 hours at 37° C. The resulting peptides were cleaned up using Sep -Pak C_18_ columns (Waters ^TM^). The peptides were then labelled with 10 plex TMT labels according to manufacturer’s protocol.

### Liquid chromatography-high resolution mass spectrometry (LC-HRMS) of proteome

The peptides were analyzed on Q Exactive HF-X Hybrid Quadrupole Orbitrap mass spectrometer (Thermo Scientific, Bremen, Germany) interfaced with Dionex Ultimate 3000 nanoflow liquid chromatography system. Peptides were separated on an analytical column 75 µm × 50 cm, RSLC C18 at a flow rate of 280 nl/ min using a gradient of 8% solvent A (0.1% formic acid in water) - 35% solvent B (0.1% formic acid in Acetonitrile) for 104 min. The total run time was kept for 120 min. The data was acquired for mass spectrometer in a data-dependent acquisition mode. The MS scan from m/z 350–1600 was acquired in the Orbitrap at a resolution of 120,000 at 200 m/z. The automatic gain control (AGC) target for MS1 was set as 4 × 10^6^ and ion filling time set at 50 ms. The most intense ions with charge state ≥ 2 was isolated and fragmented using HCD fragmentation with 34% normalized collision energy and detected at a mass resolution of 30,000 at 200 m/z. The AGC target for MS/ MS was set as 10 × 10^4^ and ion filling time set at 150 ms, while dynamic exclusion was set for 30 s with a 10-ppm mass window.

#### Proteomics data processing

The acquired mass spectrometry data were searched against Uniport *Zingiberales* database using SEQUEST HT and Mass amanda 2.0 search algorithms through Proteome Discoverer platform (version 2.1, Thermo Scientific). Following search parameters were used: two missed cleavages allowed, trypsin as cleavage enzyme, a tolerance of 10 ppm on precursors and 0.6 Daltons on the fragment ions. Fixed modification includes: carbamidomethylation at cysteine, TMT 10-plex (+229.163) modification at N-terminus of peptide and lysine and variable modification as oxidation of methionine and deamination of asparagine and glutamine. Data was also searched against a decoy database and filtered with a 1% false discovery rate (FDR). The mass spectrometry data generated from this study have been deposited to the ProteomeXchange Consortium (https://www.proteomexchange.org/) via PRIDE partner repository with the dataset identifier PXD020078.

#### Preparation of samples for metabolomics

Collar tissue samples were ground in pre-chilled mortar and pestle with liquid nitrogen to powder form. For each biological replicate 500-550 mg of the tissue were taken and extracted with 4 ml of chilled 80% LC-MS grade aqueous methanol with 0.1% of formic acid by brief sonication followed by vortexing for 5 min. Tissue extracts were then centrifuged at 16000×g for 15 min at 4°C and supernatant was collected. This supernatant was filtered in 0.22 µ nylon syringe filter (Axiva Sichem ®) and stored in 1.5 ml HPLC vials (Shimadzu) at -80 °C until analyzed.

#### Untargeted liquid chromatography-high resolution mass spectrometry (LC-HRMS) of metabolome

Supernatant from each sample were analyzed in liquid Chromatography high resolution-Mass Spectroscopy (LC-HRMS) in Orbitrap Eclipse Tribrid MS connected with UltiMate 3000 RSLC nano UHPLC or vanquish UHPLC system (Thermo Fisher Scientific, Massachusetts, United States). From a single quality control (QC) sample, multiple quality control sample aliquots were prepared. LC separation were conducted using MS grade water having 0.01% formic acid (v/v; solvent A) and methanol having 0.01% formic acid (v/v; solvent B) as the mobile phase at a flow rate of 0.35 ml/min with temperature 45⁰C. Reverse phase ACQUITY UPLC HSS T3 C_18_ column (Waters, Milford, MA, USA) of size 2.1 x 150 mm, 1.8 µm particle size (#186003540) and 100 A⁰ pore size was used. The gradient program was conducted as follows: 0 min, 0.5% B; 0-5 min, 50% B; 6 min, 98% B; 7-12 min, 98% B; 13-13.1 min, 0.5% B; 13.2-15 min, 0.5% B. The injection volume was 5 μl. The mass spectrometer was operated in positive as well as negative ion mode, and the electrospray ionization (ESI) source parameters were as follows: vaporizer temperature 400°C; ion spray voltage 3400 V (+)/2800 V (–); ion source sheath gas, auxiliary gas, and curtain gas were set at 40, 50, and 0, respectively. MS scan properties (MS OT) were set as: acquisition mode full MS/dd-MS2, resolution setting 60000, maximum injection time 50 ms and mass scan ranges at m/z 70-1000 Da. Data dependent MS2 (OT HCD) were as: resolution setting 15000, maximum injection time (ms) 22, isolation window (m/z) 1.5, Collision energies 20%, 35% and 50%, Intensity threshold 2.00E+04 and dynamic exclusion (sec) 2.5. The analysis was performed with 3 biological replicates per treatment; a total of 9 samples were analysed.

#### Metabolomics data processing

The .raw file of the mass-spectrometry data generated were processed and deconvoluted in Compound Discoverer 3.3 (Thermo Fisher Scientific^®^) by using retention time alignment, compound grouping, elemental composition prediction and gap-filling. A pre-existing workflow template titled ‘Untargeted Metabolomics using Online Databases, mzLogic, and Molecular Networks’ was used to identify the differences between samples. [M+H], [M-H], [M-H+TFA-1], [M-H-H_2_O-1] and [2M+FA-H-1] adducts were considered for mass feature (m/z) identification. Annotations of the metabolites were done by using data bases such as mzCloud, KEGG, NIST, PlantCyc and ChemSpider. The deconvoluted data was normalized with fresh weight and blank (base line peak area or noise in the extraction solvent’s chromatograms) and uploaded onto MetaboAnalyst 6.0 (Pang et al., 2024). Top 5000 features were filtered out based on interquartile range (IQR) (with 40% cut-off). The data sets were further normalized and scaled by using sum, logarithmic conversion (log_10_) and pareto-scaling and finally significantly altered (*P*<0.05) mass features were used for all univariate and multivariate analysis.

#### RNA isolation, cDNA synthesis and real time qPCR quantification of gene expression

The collar tissue samples had been collected and total RNA has been isolated from all replicates at each time point as per protocol by Salzman et al. (1999). Treatment with RNase free DNase kit (Thermofisher Scientific, Massachusetts, United States) was done to get rid of genomic DNA remnants in RNA samples. The purity of RNA was assessed using nanodrop, The cDNA preparation from 1 μg of the total RNA was done using a Verso cDNA synthesis kit (Thermofisher Scientific, Massachusetts, United States) as per the manufacturer’s protocol. The primers for respective genes were designed based on *Zingiber officinale* transcriptomic data available with investigators, and using the software Geneious v1.6.8 and were custom synthesized with Sigma (Sigma Genosys, Bengaluru, India) (Table S1). For analysis, the primers that generated a single sequence with a predicted homology in searches of databases were employed. On a Bio-Rad CFX96 real-time system, the RT-qPCR was carried out using iQ™SYBR® Green Supermix (Bio-Rad, Hercules, CA, USA). At each examined time point, four biological replicates and two technical replicates were employed. The program qbasePLUS (Biogazelle, Belgium) was used to estimate the fold change in gene expression from the Cq values. In comparison with the 0 hpi and after normalizing with the reference genes actin and eukaryotic elongation factor (eEF), the fold change of a gene at various time points following pathogen inoculation was calculated.

In case of *P. myriotylum* gene expression estimation, RT-qPCR was performed after total RNA extraction using Trizol and cDNA preparation. Expressions were profiled for four pathogenicity associated genes i.e. *cellulase*, two *glycoside hydrolases* (*GH1a, GH3*) and *necrosis inducing proteins* (*NIP1*). These genes where checked for their relative expression in different concentrations of TGA treatments such as 0.1 mM and 0.3 mM against the control (1% DMSO). At each treatment, four biological replicates and two technical replicates were employed. In comparison with the control ‘cq values’ and after normalizing with the reference housekeeping genes actin and beta tubulin, the fold change of a gene expression at various treatments were calculated. All the primers are listed in supplementary table S1.

#### Statistical and bioinformatics analysis

All the univariate and multivariate statistical analysis was conducted by using MetaboAnalyst 6.0 (https://www.metaboanalyst.ca/), Heatmapper (http://heatmapper.ca) and GraphPad Prism (version 10.0) (https://www.graphpad.com). Relative distance plasticity index (RDPI) was calculated as described by Valladares et al., 2006 and Li et al., 2020. Bioinformatics analysis with proteomics data was conducted by using UniProt (https://www.uniprot.org/), Shiny GO (Version 0.82) (https://bioinformatics.sdstate.edu/go/) and STRING (https://string-db.org/). For metabolomics data based bioinformatics we used MBROLE (version 3.0) and MetaboAnalyst 6.0 (https://www.metaboanalyst.ca/). Classifier identification through prediction of variable importance was done through two machine learning models: Random Forest (RF) and Support Vector Machine (SVM). Validation through receiver operating characteristic curve (ROC) was performed by using GraphPad Prism. All the analysis was conducted by using significantly altered proteome and metabolome considering control-12 hpi and control-24 hpi. Significance differences for multi-set data were calculated by one-way ANOVA followed by Tukey’s post-hoc test and for two set data unpaired *t*-test were performed.

#### TGA agar diffusion assay

The *trans*-geranic acid (TGA) was tested to check its antifungal property against oomycete *Pythium myriotylum* using agar diffusion assay. Stock solutions of each testing solution of concentrations 0.1 mM, 0.3 mM, 1 mM and 5 mM was prepared by dissolving the compound in 1% DMSO (v/v), according to its molarity. Using a sterile cork borer, punched two wells in each sides of Potato Dextrose Agar (PDA) plate, approximately equidistant from the center and each other. TGA solution was added to well in 100 µl volume of respective concentration. Control well was filled with the same volume 100 µl of 1% DMSO. Each concentration was tested with minimum three replicates. Each plate was directly inoculated the in center of the two wells of PDA plates with a mycelial agar plug (10 mm in diameter) from the edge of an actively growing 48 hour old culture. All the plates were incubated at 25– 28°C for 48–72 hours to observe the zone of inhibition.

#### 2.12 TGA suspension culture assay

To check the antifungal activity of TGA in suspension culture, different stock concentrations such as 0.1 mM, 0.3 mM, 1 mM and 5 mM was prepared. To 100 ml Potato dextrose broth (PDB-2.4%), respective concentrations were added each with 3 replicates. Two mycelial agar plugs from the periphery of an actively growing *P. myriotylum* solid culture was inoculated and the suspension cultures were incubated at 28°C in a shaking incubator in the dark, with 150 rpm shaking for 6 days to check the bioactivity of TGA. After 6 days, visualization was done. Further, cultures were filtered, squeezed, dabbed in filter paper to remove excess PDB media and weighed for each treatment. A minimum of three biological replicates were taken for each treatment.

### Sequencing confirmation of *P. myriotylum*

The *P. myriotylum* samples collected from suspension culture assay was checked to confirm the presence of *Pythium*. Total DNA was isolated from the samples using Nucleospin isolation kit (Machery-Nagel, Düren, Germany) according to the manufacturer’s protocol. The purity of DNA was assessed using nanodrop. PCR confirmation was done using ITS primers and sample was submitted for Sanger sequencing. Confirmation of *P. myriotylum* sequence was done using BLAST.

### Necrotic lesion study in model plant *Nicotiana benthamiana*

A 48 hour old *P. myriotylum* mycelial disc from PDA (3 mm diameter) was inoculated onto the leaves of 4 week old *N*. *benthamiana* plants at four-leaf stage. A solution of 1 mM TGA in 1% DMSO (v/v), was sprayed on the discs immediately after inoculation. The control plants were sprayed with 1% DMSO. The size (in mm^2^) of necrotic lesions were measured (mean ± standard deviation) at 5 dpi. Twelve replicates for each group (TGA and control) were used in each experiment.

## Results

### Optimized integrative omics pipeline for deciphering defense metabolism

*P. myriotylum* colonization in the collar region of ginger was confirmed by trypan blue staining based microscopy (Figure 1) and based on that we performed a temporal pre-optimized mass-spectrometry based proteomics and metabolomics (Figure S1a). Thereafter, we applied an integrative computational data analysis pipeline for obtaining a comprehensive defense metabolic map of ginger against *P. myriotylum* (Figure 1). Integrity of the biological replicates was checked by using proteomics data-sets analyzed with correlation matrix and partial least square discriminant analysis (PLS-DA) model which showed clear differentiation among the sample sets (biological replicates from each treatment i.e. control, 12 hpi and 24 hpi) (Figure S1b, c). To check the correlation of the proteome and metabolome alteration we performed a Pearson’s correlation analysis and observed a high correlation at 12 hpi (PCC = 0.9621) and 24 hpi (PCC = 0.9398) compared to control (Figure S2a, b). This result suggests host’s alterations of proteome and metabolome in response to *P. myriotylum* infection are strongly correlated.

**Fig. 1.**
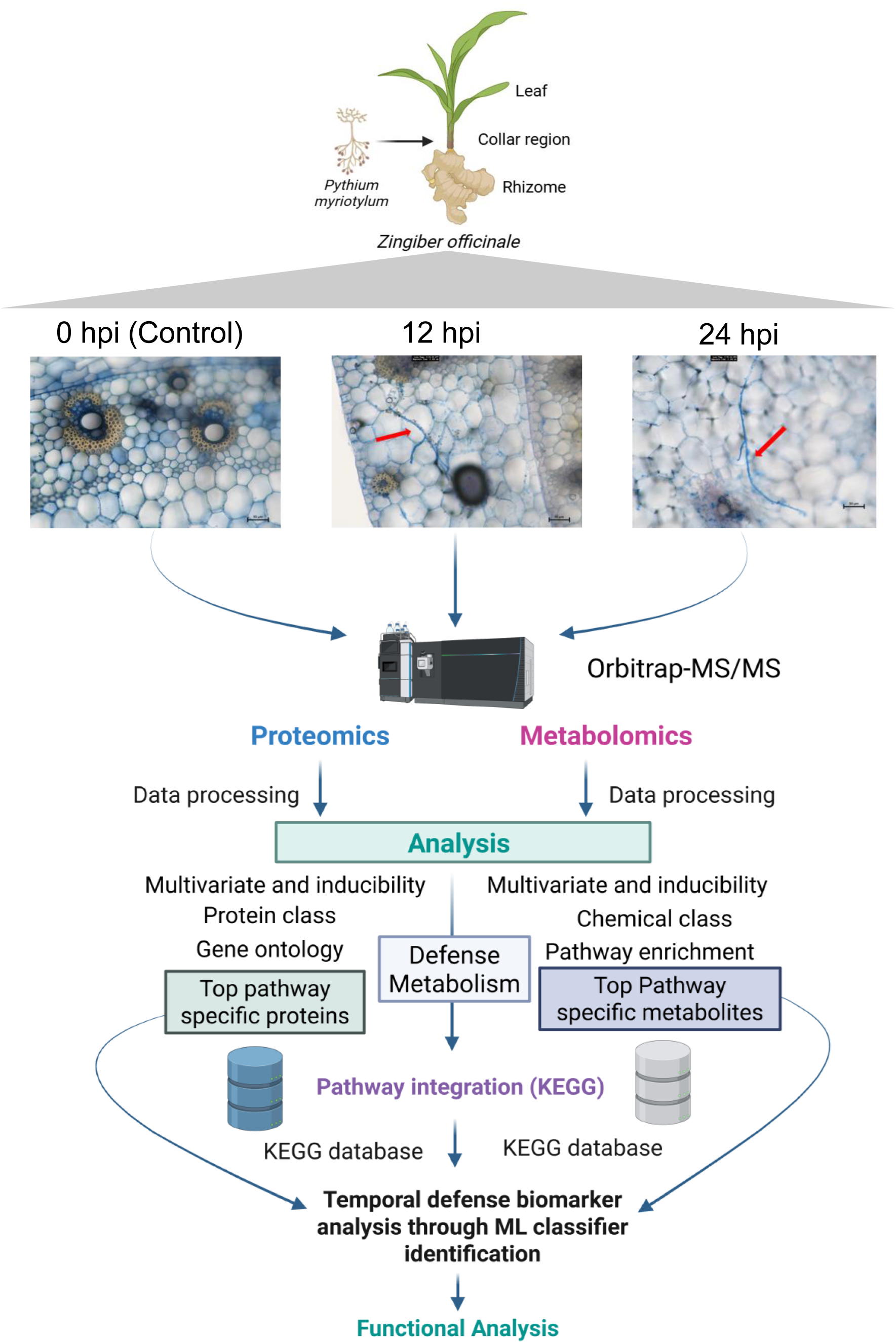
Overview of work flow and analysis pipeline. *Pythium myriotylum* infected ginger (collar region) tissues were collected at three different time points (0 hpi, 12 hpi and 24 hpi) and were analyzed for mass spectrometry based proteomics and metabolomics. Both the ‘omics’ data were processed and analyzed with optimized computational pipeline and integrated. Identification of classifiers was done by classification machine learning models (i.e. random forest, SVM) based approach. Finally identified best classifier was functionally validated. The images were created in BioRender. (https://BioRender.com/)

### Proteomics revealed cellular detoxification and secondary metabolism are crucial defense responses against *P. myriotylum*

We observed 1140 and 1095 proteins were significantly (*P*<0.05) altered at 12 hpi and 24 hpi respectively (Figure 2a; Table S2; Table S3) and among them 799 proteins were commonly altered (Figure 2b, Table S4). Further, to measure inducibility of the whole proteome in response to pathogen infection, we computed Relative Distance Plasticity Index (RDPI) and observed that proteome at 12 hpi showed significantly high inducibility (*P*<0.05) compared to 24 hpi (Figure 2c). From the clustered (Pearson’s correlation linkage on rows) heat map we observed at 12 hpi 461 proteins were up-regulated and 679 proteins were down-regulated (Figure 2d) while at 24 hpi 618 proteins are up-regulated and 477 proteins are down-regulated (Figure 2e). Gene Ontology (GO) analysis based on ‘biological activity’ showed 268 and 249 pathways were significantly (FDR<0.05) enriched at 12 hpi and 24 hpi respectively (Figure 2f; Table S5). Among these enriched pathways top five pathways were observed to be associated with cellular detoxification process (Figure 2g; Table S6). Further, Kyoto Encyclopedia of Genes and Genomes (KEGG) pathway enrichment analysis (Figure. S3a, b) with dysregulated proteins showed four pathways common to 12 hpi and 24 hpi as top pathways. Among these pathways ‘Biosynthesis of secondary metabolites’ is directly associated with defense mechanism (Figure 2h). Interestingly, protein class enrichment analysis showed higher number of ‘metabolites interconversion enzymes’ at 12 hpi compared to 24 hpi which supports the higher inducibility of proteome at 12 hpi by indicating synthesis of pathway enzymes (Figure. S3c). Next, to check whether detoxification proteins and proteins associated with secondary metabolite biosynthesis work together, we mapped these common proteins that are altered at 12 hpi and 24 hpi onto protein-protein interaction network and observed strong interactions (PPI enrichment *P*<1.0e^-16^) of cellular detoxifying proteins with proteins of secondary metabolites biosynthesis (Figure 2i) indicating co-regulation of cellular detoxification process with secondary metabolism upon *P. myriotylum* colonization. By comparing with UniProt database we identified 47 cellular detoxification associated proteins and by analysis with a Pearson’s correlation based clustered heat map we observed that 12 hpi and 24 hpi datasets are closely linked indicating high correlation of their alteration patterns (Figure 3a). Though, some of these detoxifying proteins were down-regulated; most of them were highly up-regulated either at 12 hpi or 24 hpi suggesting their involvement in defense responses against *P. myriotylum* at both the early and late phase of defense.

**Fig. 2.**
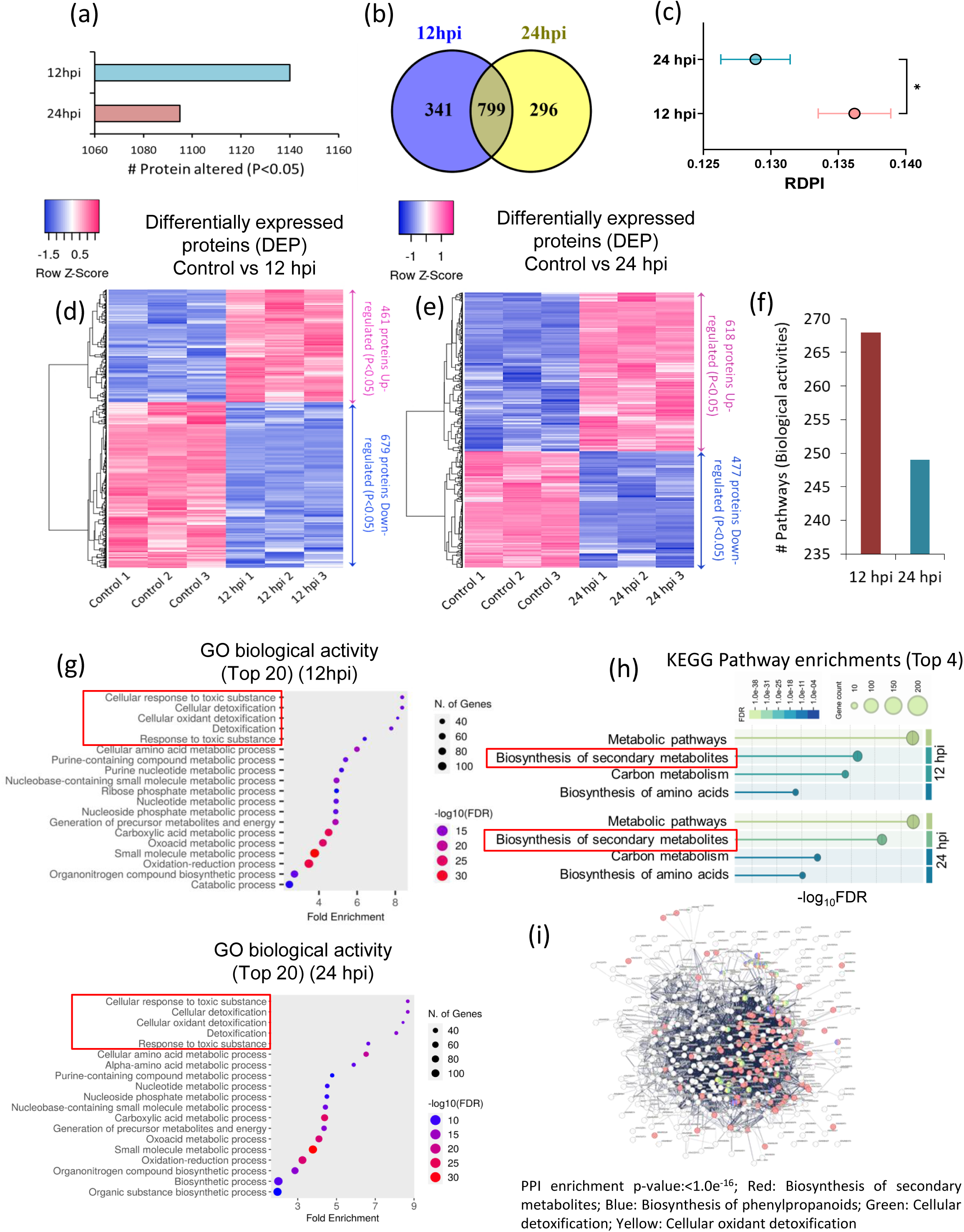
Differentially altered proteome in *Pythium myriotylum* infected ginger (collar region) tissues at three different time points (0 hpi, 12 hpi and 24 hpi). (a) Number of proteins significantly (*P*<0.05) altered at 12 hpi and 24 hpi compared to control (0 dpi). (b) Venn diagram indicates numbers of specific and common proteins significantly altered at 12 hpi and 24 hpi. (c) Inducibility in terms of relative distance plasticity index (RDPI) of significantly altered proteins at 12 hpi and 24 hpi. Hierarchically clustered heat map (Pearson’s correlation, complete linkage) of differentially altered (*P*<0.05) proteins in (d) control Vs 12 hpi and (e) control Vs 24 hpi. (f) Number of gene ontology (GO) metabolisms enriched (FDR<0.05) in *Z. officinale* after 12 hpi and 24 hpi. (g) Top 20 gene ontology (GO) metabolisms based on fold enrichment. Red squares indicate top five metabolisms in each treatment. (h) Top four metabolisms enriched with GO analysis with KEGG metabolic pathway enrichment analysis at 12 hpi and 24 hpi Red squares indicate defense associated ‘Biosynthesis of secondary metabolites’ pathway. (i) Protein-protein interaction network analysis (enrichment *P*<1.0e^-16^) shows the a sub-set of proteins associated with the cellular detoxification and secondary metabolism interact. Colours of nodes indicates protein’s co-associations with detoxification and secondary metabolism.

**Fig. 3.**
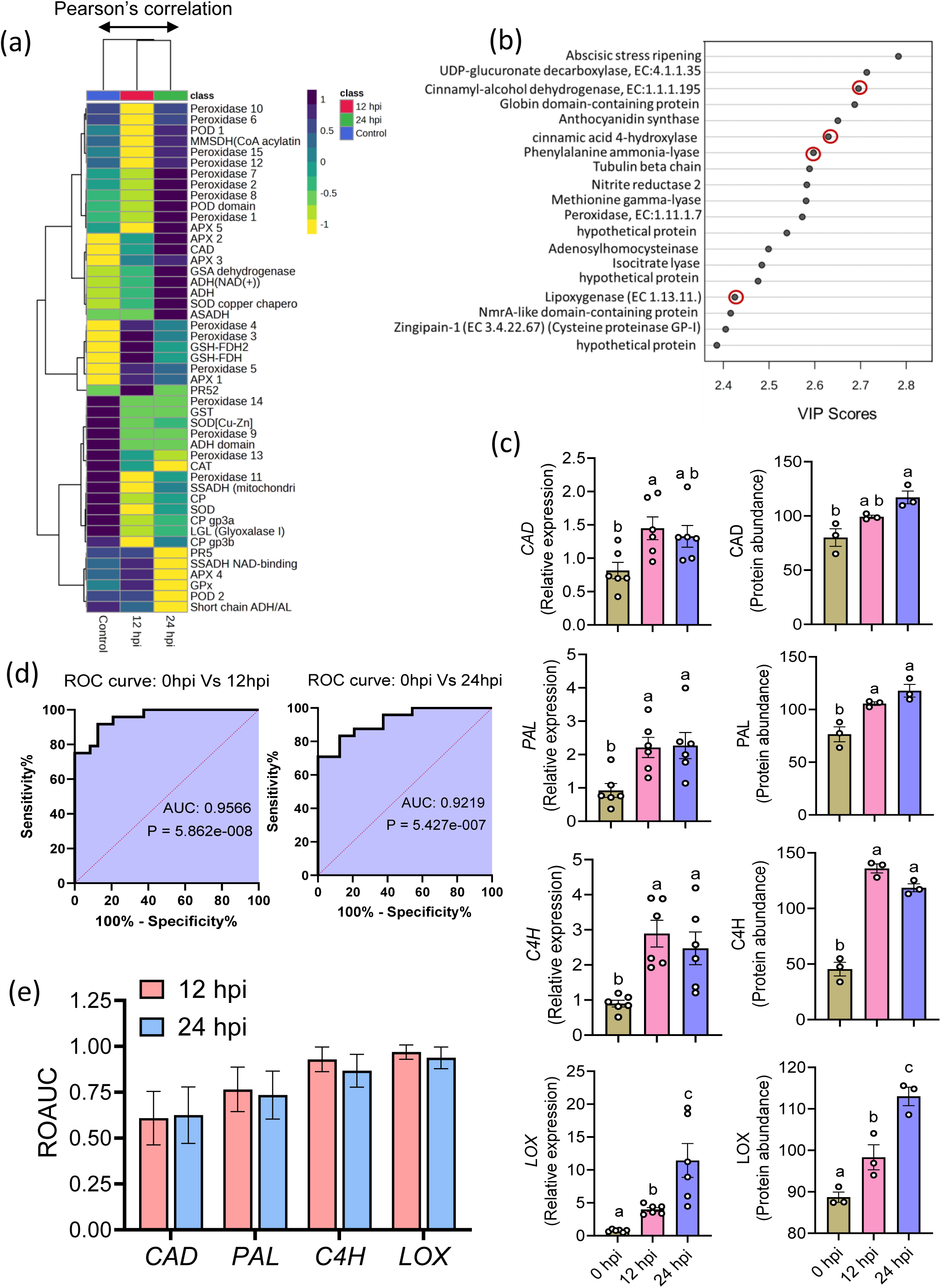
Identification of detoxification and secondary metabolism associated proteins in the collar region of *Pythium myriotylum* infected ginger at three different time points (0 hpi, 12 hpi and 24 hpi). (a) Hierarchically clustered (Pearson’s correlation) heat map of top five metabolism associated 47 detoxifying ginger proteins at three different time points after infection with *P. myriotylum*. (b) VIP score plot of top 19 proteins extracted from PLS-DA classification model among 0 hpi, 12 hpi and 24 hpi proteome data sets. Red circles indicate major proteins associated with defense metabolism. (c) Comparative mean ± SE of relative expressions and protein abundances of cinnamyl-alcohol dehydrogenase (CAD), Phenylalanine ammonia-lyase (PAL), cinnamic acid-4-hydroxylase (C4H) and lipooxygenase (LOX) in ginger after 0 hpi, 12 hpi and 24 hpi. Significant differences were calculated by one way ANOVA followed by Tukey’s post-hoc test. (d) Multivariate receiver operating characteristic curve (ROC) analysis of the four selected proteins (gene expressions) at 12 hpi and 24 hpi compared to control. (e) Univariate ROAUC ± SE of CAD, PAL, C4H and LOX proteins at 12 hpi and 24 hpi.

### Jasmonic acid and phenylpropanoid biosynthetic proteins are key defense associated proteome classifiers of *P. myriotylum* infection

To identify the subset of classifier proteins dominating the defense-associated proteome alteration, we used a classification PLS-DA machine learning model (Figure S1c) to compute the Variable Importance in Projection (VIP) scores. VIP score plot of top 19 proteins indicated four proteins directly associated with phenylpropanoid biosynthesis: cinnamyl-alcohol dehydrogenase (CAD), anthocyanidin synthase (ANS), cinnamic acid 4-hydroxylase (C4H), phenylalanine ammonia-lyase (PAL) and one protein, lipooxyginase (LOX), is associated with jasmonic acid biosynthesis (Figure 3b). To validate these five proteins as classifiers we checked the level of expressions of the genes of these proteins and compared with the corresponding protein abundances (Figure 3c, Figure S4a, b). Except ANS, all the other proteins (CAD, C4H, PAL and LOX) showed significant up-regulation at 12 hpi and 24 hpi and their gene expressions synchronized with the protein abundances. Therefore, we used the gene expression data of these four proteins to cross-validate them to be classifiers with multivariate ROC curve analysis. High area under the curve (AUC) values were observed for both the 12 hpi and 24 hpi datasets compared to control which were AUC = 0.9566 and AUC = 0.9219 respectively (Figure 3d). Additionally, *PAL*, *C4H* and *LOX* genes showed high univariate AUC predicting them as highly potential classifiers involved in defense (Figure 3e). We also found that alteration of LOX, the jasmonic acid (JA) biosynthetic enzyme, has strong correlations (Minkowski) with *PAL* and *CAD* (Figure S4c) indicating jasmonic acid mediated modulation of secondary metabolism.

### Metabolomics confirmed specific secondary metabolic dysregulation as major defense responses against *P. myriotylum*

Next, through high resolution untargeted metabolomics study we deconvoluted 12255 metabolite features (Table S7) and observed *P. myriotylum* infection has great impact on the overall metabolome plasticity. Clustered heat map (Pearson’s correlation on row) showed most distinct metabolome alteration pattern at 24 hpi (Figure S5a). A cross validated PLS-DA classification model showed clearly separated sample clusters of control, 12 hpi and 24 hpi (Figure S5b, c). In the whole metabolome, 379 and 828 metabolite features (m/z) were significantly (*P*<0.05) dysregulated at 12 hpi and 24 hpi respectively, compared to control where 98 metabolites are commonly dysregulated at these two time points (Figure 4a, b; Table S8). RDPI of the whole metabolome in response to pathogen infection indicated significant higher inducibility (*P*<0.05) at 24 hpi compared to 12 hpi (Figure 4c). Pearson’s correlation based clustered heat maps showed 86 up-regulated and 293 down-regulated features at 12 hpi while 529 up-regulated and 299 down-regulated features at 24 hpi (Figure 4d, e). To check the metabolic alterations based on metabolomics data sets, we performed chemical-class enrichment (Figure S6) based identification of KEGG ids of the significantly altered metabolite features at 12 hpi and 24 hpi. These KEGG ids were used for pathway enrichment analysis on the KEGG pathway map separately for these two time points. Only significantly enriched pathways (FDR<0.05) were considered and top 20 pathways were plotted (Figure 4f, g). Among these pathways, five pathways at 12 hpi and seven pathways at 24 hpi were enriched those are directly associated with secondary metabolism. This result indicates secondary metabolic pathways in ginger are crucially triggered as defense response against *P. myriotylum* in both the early and late phase of defense.

**Fig. 4.**
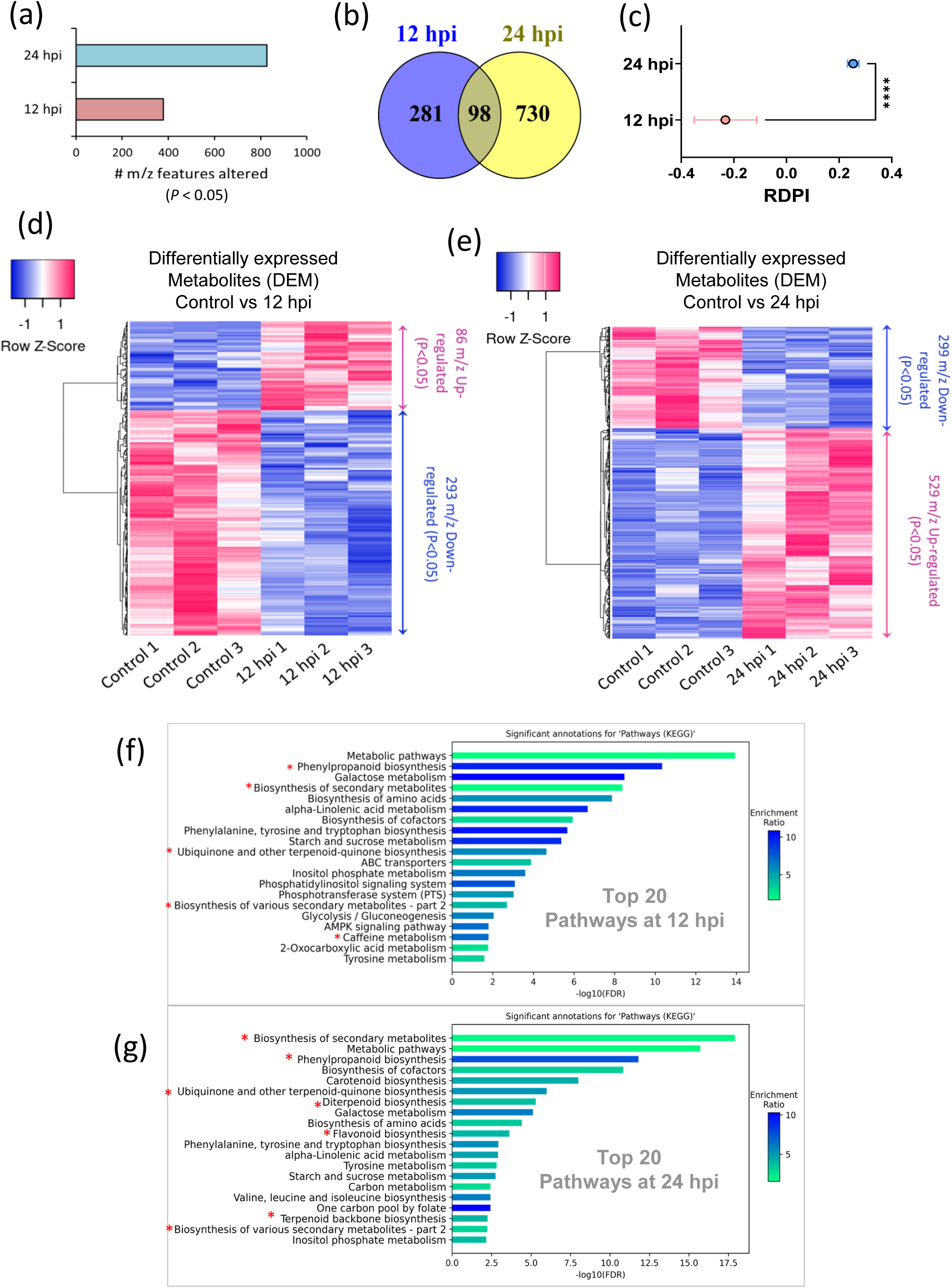
Differentially altered metabolome in *Pythium myriotylum* infected ginger (collar region) tissues at three different time points (0 hpi, 12 hpi and 24 hpi).. (a) Number of metabolite features significantly (*P*<0.05) altered at 12 hpi and 24 hpi compared to control (0 hpi). (b) Venn diagram indicates numbers of specific and common metabolite features significantly altered at 12 hpi and 24 hpi. (c) Inducibility in terms of relative distance plasticity index (RDPI) of significantly altered (*P*<0.05) metabolite features at 12 hpi and 24 hpi. Hierarchically clustered heat map (Pearson’s correlation) of differentially altered (*P*<0.05) metabolite features in (d) control Vs 12 hpi and (e) control Vs 24 hpi. KEGG metabolisms enriched (FDR<0.05) in *Z. officinale* after (f) 12 hpi and (g) 24 hpi.

### Machine learning models predicted *trans*-geranic acid (TGA) as top defense classifier metabolite

By integration of proteomics and metabolomics based pathway enrichment data through Venn diagram we identified 10 metabolic pathways those are ‘commonly enriched’ at both 12 hpi and 24 hpi (Figure 5a) 4 among them are defense associated. The reason behind considering the ‘commonly enriched’ pathways was the presence of maximum specialized or secondary metabolisms in the common pathway group. We annotated 23 specialized metabolites of these 4 defense metabolic pathways (Table S9). Pearson’s correlation based clustered heat map indicated that most of the metabolites are significantly up-regulated at 24 hpi (Figure 5b). RDPI assessment showed significantly (*P*<0.05) higher inducibility of these 23 specialized metabolites set at 24 hpi compared to 12 hpi suggesting these metabolites were induced mostly at the late phase of infection. Gingerdione, gingerol, shogaol, curcumin are known defensive bioactive specialized metabolites in ginger and caffeoyl-coenzyme A O-methyl transferase (CCOMT) is one of the major biosynthetic enzymes for these metabolites (Li et al., 2021). We observed that the level of these 4 metabolites and expression of *CCOMT* gene were significantly (*P*<0.05) up-regulated at 24 hpi (Figure S7). To identify the most crucial metabolites involved in defence, we applied two machine learning models on the data set of 23 specialized metabolites; those are random forest (RF) and support vector machine (SVM). We plotted top 5 metabolites predicted by these models at both 12 hpi and 24 hpi. Based on ‘average importance’ of the metabolites (features) *trans*-geranic acid (TGA) was found to be the top feature in three models (12 hpi random forest, 12 hpi SVM and 24 hpi random forest) (Figure. 5d,e,f) while jasmonic acid (JA) was the top feature in 24 hpi SVM model and TGA was second top (Figure 5g). TGA was also observed as mostly up-regulated metabolite in the volcano-plot analysis with control and 24 hpi metabolomics data (Figure 5h). Both JA and TGA were found to be up-regulated at both 12 hpi and 24 hpi and showed positive correlation (Figure 5i; Figure S8). For confirming the induction of TGA biosynthesis upon *P. myriotylum* infection, we checked the expression of a key TGA biosynthetic pathway enzyme, geraniol dehydrogenase (GeDH) at both protein and transcript level and observed significant inductions at 24 hpi (Figure 5j). This result suggests TGA biosynthesis is crucial as a defense response against *P. myriotylum*.

**Fig. 5.**
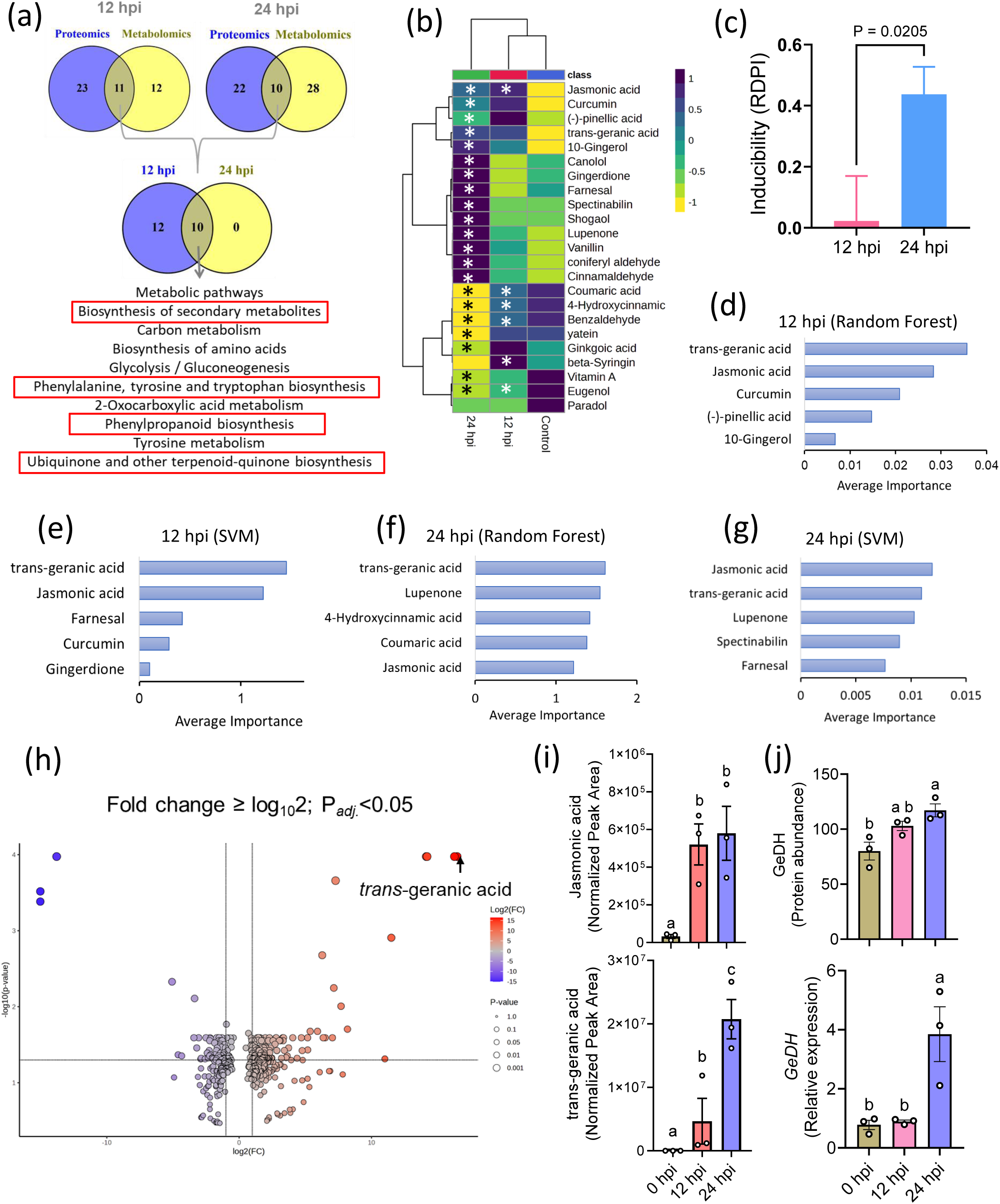
Integration of major defense pathways enriched in proteomics and metabolomics data with identification of major metabolites involved in defense against *P. myriotylum*. (a) Venn diagram mediated identification of ten common metabolic pathways enriched in both proteomics and metabolomics data at 12 hpi and 24 hpi. (b) Hierarchically clustered heat map (Pearson’s correlation, complete linkage) of 23 annotated metabolites associated with defense associated 4 common metabolic pathways. * indicates significance difference (*P*<0.05) of 12 hpi and 24 hpi with the control. (c) Inducibility in terms of RDPI of 23 annotated metabolites at 12 hpi and 24 hpi. Statistical significance was analyzed through unpaired *t*-test (*P* < 0.05 was taken as cut-off). Testing of machine learning models to find out top five feature of importance by (d) random forest at 12 hpi, (e) SVM at 12 hpi, (g) random forest at 24 hpi, SVM at 24 hpi. (h) Volcano plot analysis (Log_2_ Fold Change ≥ 2) of significantly altered (*P*<0.05) metabolites at 24 hpi indicating *trans*-geranic acid as mostly up-regulated metabolite. (i) Mean ± SE of normalized peak area of jasmonic acid and trans-geranic acid at three different time points. (j) Mean ± SE of protein abundances and relative expressions of geraniol dehydrogenase (GeDH) at three different time points. Statistical significance was analyzed through one way ANOVA with Tukey’s Post-hoc test. Different letters indicate significant differences (*P*<0.05 was taken as cut-off).

### TGA showed strong bioactivity against *P. myriotylum*

For functional analysis of TGA, we checked the effect of TGA in different concentrations on the *in vitro* grown *P. myriotylum*. We used 0.1, 0.3, 1 and 5 mM concentrations on both the solid (in PDA) and suspension (in PDB) culture of *P. myriotylum*. First, we confirmed the purity of culture grown for treatment by Sanger sequencing and then performed an agar diffusion assay. We observed that inhibition of pathogen growth initiated with 0.1 mM followed by maximum inhibition with 5 mM TGA (Figure 6a). For quantitative inhibition assessment we performed suspension culture assay and measured the fresh weight of the grown mycelia after 6 days of incubation. In this assay, a stronger inhibition was observed where 0.1 mM TGA showed 78.25% inhibition followed by increased to complete inhibition of growth with 0.3 mM (87.9% inhibition), 1 mM (100% inhibition) and 5 mM (100% inhibition) respectively (Figure 6b, c). This result suggests TGA is strongly active against *P. myriotylum* growth. Next, to see whether TGA has any effect on pathogenicity of *P. myriotylum* we assessed the expression levels of 4 pathogenicity associated genes i.e. *NIP1* (*Necrosis inducing protein1*), *GH3* (*Glycoside Hydrolase 3*) and *GH1* (*Glycoside Hydrolase 1*), in TGA treated *Pythium* mycelia. To avoid concentration bias we used two low concentrations of TGA: 0.1 mM and 0.3 mM, for this experiment and observed both the concentrations of TGA significantly reduced the expression of *NIP1* and *GH1*, while 0.3 mM reduced the expression of *GH3* and cellulase (Figure 6d). This result suggests TGA reduces pathogenicity of *P. myriotylum*. We further checked *in plant* bioactivity of TGA against *P. myriotylum* growth. For this analysis we used *Nicotiana benthamiana* leaves for convenience in lesion size measurement. We inoculated *N. benthamiana* leaves with *P. myriotylum* and immediately sprayed 1mM TGA solution (in 0.5% DMSO) while control leaves were sprayed with 0.5% DMSO. We observed significantly smaller lesion sizes in TGA treated leaves after 5 days of inoculation indicating protective nature of TGA against *P. myriotylum* (Figure 6e, f).

**Fig. 6.**
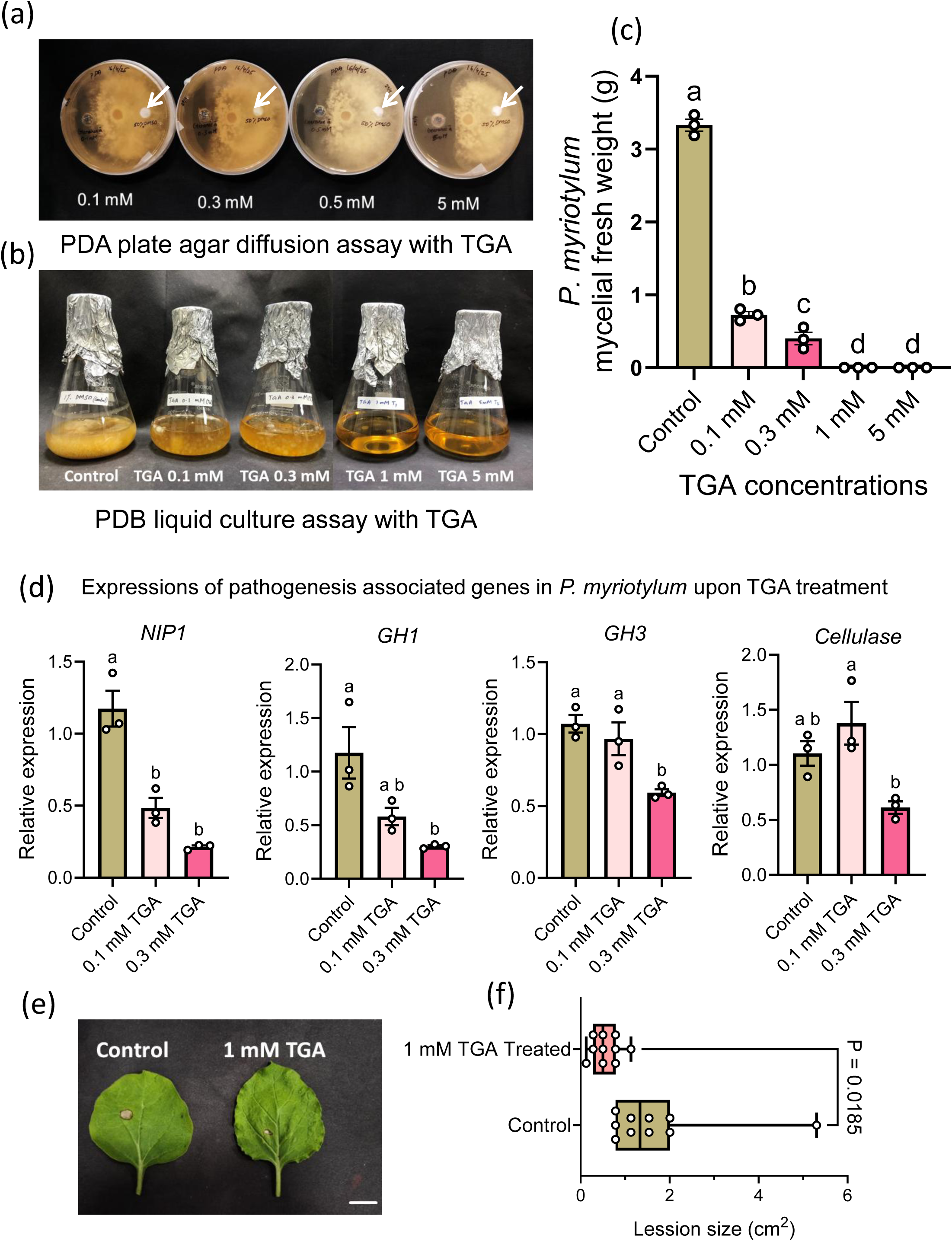
Bioactivity of *trans*-geranic acid (TGA) as anti *P. myriotylum* defense metabolite. (a) Agar diffusion growth assay of *P. myriotylum* culture with different TGA concentrations. Arrows indicate control. (b) growth assay of *P. myriotylum* suspension culture in PDB with different TGA concentrations. (c) Mean ± SE of *P. myriotylum* mycelial fresh weight in liquid culture after treatment with different TGA concentrations. (d) Mean ± SE of expressions of pathogenesis associated genes in *P. myriotylum. NIP1* (*Necrosis inducing protein1*), *GH3* (*Glycoside Hydrolase 3*) and *GH1* (*Glycoside Hydrolase 1*). Statistical significance was analyzed through one way ANOVA with Tukey’s Post-hoc test. Different letters indicate significant differences (*P*<0.05 was taken as cut-off). (e) Representative images of *in planta* lesion size of *P. myriotylum* infection on *Nicotiana benthamiana* leaves. Scale represents 1 cm. (f) Box-whisker plots (minimum to maximum values) of lesion area in control and TGA treated leaves (n =10). Statistical significance was analyzed through unpaired *t*-test (*P* < 0.05 was taken as cut-off).

## Discussions

Worldwide ginger production has been adversely affected by various pathogens such as *Pythium* species which cause rhizome rot (Webster and Weber, 2007), *Fusarium oxysporum* causing yellow disease (Simmonds, 1956), *Phyllosticta zingiberi* triggering phyllosticta leaf spot disease (Ramakrishnan, 1942), *Ralstonia solanacearum* causing bacterial wilt (Yabuuchi et al., 1995). Amongst these, rhizome rot is the deadliest disease which poses serious risk to global annual ginger production (Dohroo, 2005) and can devastate fields, sometimes approaching complete yield loss. Therefore, understanding defense metabolism in ginger against the soft-rot oomycete *Pythium myriotylum* is pivotal for resilient cultivation and targeted crop protection.

Here, we employed an integrative quantitative proteomics and untargeted metabolomics strategies to investigate the temporal dysregulation of metabolisms associated with key defense sectors in ginger upon *P. myriotylum* infection. Quantitative proteomics through TMT labelling with no missing values having advantage over data dependent acquisition (DDA) label-free proteomics approaches (Song et al., 2025) that provides precise information regarding protein level alterations with identification of biomarkers in plants (Luan et al., 2022). Therefore, we applied this approach and through gene ontology analysis we observed a significantly high inducibility of proteome at 12 hpi compared to 24 hpi which suggests at early phase of infection biosynthesis and modifications of proteins happened more rigorously for helping the plants to produce enzymes for driving the metabolic pathways which biosynthesize metabolites at late phase. This data was confirmed by protein class enrichment analysis where ‘metabolite interconversion enzymes’ were higher in number at 12 hpi compared to 24 hpi. Eventually, biological activity based GO analysis with annotated proteins exhibited cellular detoxification metabolisms as top enriched pathways where 47 cellular detoxification or reactive oxygen species (ROS) associated proteins were involved. ROS production is a crucial hypersensitive immune response in plant hosts. When plants encounter pathogens, they rapidly generate ROS such as superoxide anion (O_2_^-^), hydrogen peroxide (H_2_O_2_), and hydroxyl radicals as parts of the oxidative burst that contributes to defense signaling and hypersensitive response. Excessive ROS causes cellular damage by oxidizing lipids, proteins, and nucleic acids. Thus, efficient detoxification systems are required to maintain redox homeostasis while still allowing ROS to act as defense signals (Apel & Hirt, 2004). Detoxification also requires the activity of cytochrome P450s, glutathione-S-transferases (GSTs), and ABC transporters, which metabolize and compartmentalize pathogen-derived toxins and oxidative byproducts (Edreva, 2005). Therefore, enriched detoxification pathways indicate initiation of ROS mediated defense in the infected tissue. We observed up-regulation of higher number of detoxifying proteins at 24 hpi compared to 12 hpi. However, down-regulation of some detoxifying proteins upon infection was also observed. Plants with impaired detoxification pathways often exhibit hypersensitivity or susceptibility to pathogens, whereas priming of antioxidant and detoxification systems enhances disease tolerance (Mittler, 2017; Sharma et al., 2012). Therefore, down regulation of specific detoxification proteins might be the factor behind susceptibility of *Z. officinale* to *P. myriotylum*.

Protein class enrichments showed maximum numbers of proteins were ‘metabolite interconversion enzymes’ which are generally the key driving factors of metabolic pathways. KEGG pathway enrichment analysis with proteome data showed ‘biosynthesis of secondary metabolites’ is the second of top 4 four enriched pathways and directly involved in plant defense (Erb and Klebstein, 2020). For understanding the simultaneous participations of the detoxifying proteins and secondary metabolic pathway proteins in defense, a protein-protein interaction network was created and it was observed that these proteins have strong interactions and some of them are co-involved in both cellular detoxification and secondary metabolite biosynthesis. The correlation between plant secondary metabolism and cellular detoxification during ROS production is tightly linked, as secondary metabolites often function as both antioxidants and modulators of defense signaling. For instance, phenylpropanoids and flavonoids act as ROS scavengers, directly neutralizing singlet oxygen, hydroxyl radicals, and hydrogen peroxide while stabilizing redox balance (Agati et al., 2012). Importantly, the up-regulation of secondary metabolic pathways is often ROS-dependent.

ROS act as signals to activate transcription factors (e.g., MYB, bHLH, WRKY), which induce phenylpropanoid, flavonoid, and terpenoid biosynthesis genes. This creates a feedback loop in which ROS induce secondary metabolites, and these metabolites, in turn, detoxify excess ROS, maintaining homeostasis while sustaining defense (Sharma et al., 2012). Among the identified potential classifiers proteins of *P. myriotylum* infection CAD, C4H and PAL are crucial enzymes in phenylpropanoid pathway while LOX is the biosynthetic enzyme of JA anabolism (Wasternack and Song, 2016). JA signaling strongly induces the phenylpropanoid pathway, enhancing the biosynthesis of flavonoids, lignin, and phenolic acids, which contribute to pathogen resistance and oxidative stress detoxification (Sohn et al., 2022). Therefore, in our results, significant up-regulation of all these proteins suggests activation of JA mediated defense through phenylpropanoids upon *Pythium* infection.

Inducibility of the overall metabolome at 24 hpi was found to be higher compared to 12 hpi. This showed a reverse pattern of proteome inducibility which signifies that the induced proteome dominated by ‘metabolite interconversion enzymes’ at 12 hpi resulted in the induced biosynthesis of the metabolite levels at 24 hpi. KEGG pathway enrichment with metabolome data showed various secondary metabolic pathways including phenylpropanoids, flavonoids and terpenoids biosynthesis to be enriched, in particular, at 24 hpi confirming the proteomics based pathway enrichment results. Annotations of these pathway-associated 23 metabolites were from the specialized/secondary metabolite class. Gingerol, shogaol, curcumene, gingerdione were previously reported as major anti-microbial metabolites (Gunasena et al., 2022, Beristain-Bauza et al., 2019) and CCOMT was observed as one of the major regulatory enzyme in biosynthesis of these metabolites (Li et al., 2021). We observed a significant up-regulation of these four metabolites as well as expression level of *CCOMT* gene and protein at 24 hpi, which indicate these metabolites are also participating in defense response upon *P. myriotylum* infection.

Interestingly, classification based machine learning models (i.e. random forest and SVM) did not predict gingerol, shogaol, curcumene, or gingerdione as top classifier, instead TGA, an open-chain monoterpenoid was predicted as the top hit in term of ‘average importance’ in both the models indicating it as a potential defense metabolite. Further *in-vitro* functional assays with different concentrations of TGA showed strong inhibition of *P. myriotylum* growth even with a low concentration such 0.1 mM and 0.3 mM while 1 mM and 5 mM showed complete inhibition of mycelia growth. This result confirmed TGA as a strong anti-*P. myriotylum* metabolite. TGA was previously reported to have anti-fungal activity against

*Colletotrichum graminicola* and *Fusarium graminearum* (Becher D et al, 2014), but not against *Pythium* or other oomycete. Moreover, we also found this molecule works against pathogenicity of *P. myriotylum* as expressions of four of the major pathogenicity-associated genes including genes of cell wall degrading enzymes (i.e. *NIP1*, *GH3*, *GH1* and cellulase) were supressed with TGA. The *NIP1* gene in oomycete encodes for Nep1-Like Proteins (NLPs), which are important pathogenicity factors secreted by oomycetes like *Pythium* species to induce cell death in plants (Yang et al., 2022). *GH* (Glycoside Hydrolase) genes play crucial roles in encoding enzymes that break down complex carbohydrates, which are essential for the oomycete’s growth, development, and ability to infect plants (Shen et al., 2020). On the other hand, cellulase enzyme functions for the oomycete’s pathogenicity, facilitating hyphal penetration of plant cell walls by breaking down cellulose (Zerillo et al., 2013). Therefore, suppression of these genes confirmed that TGA acts against the pathogenicity of *P. myriotylum*. Finally *in planta* functional assay showed significantly less infection in TGA treated *N. benthamina* leaves indicating the protective nature of TGA by showing bioactivity against *P. myriotylum*.

In conclusion, our integrative proteomics and metabolomics study has shown *P. myriotylum* infection induces cellular detoxifying proteins in association with secondary metabolic pathway enzymes as key defense responses in ginger. At early phase of infection ‘metabolite interconversion enzymes’ are synthesized which further get involve in biosynthesis of secondary metabolites. Secondary metabolisms, in particular, phenylpropanoids and terpenoid biosynthesis is crucial for developing defense against *P. myriotylum*. JA biosynthesis is positively correlated and alters the levels of top defense-associated classifier proteins of phenylpropanoid pathways. Moreover, a monoterpene, TGA was identified as top classifier metabolite which is induced upon infection and strongly functions *in vitro* against *P. myriotylum* by inhibiting its growth and pathogenicity linked genes. Therefore, this study has facilitated our understanding of defense metabolic trajectories in edible ginger against *P. myriotylum* and established TGA as a potential bio-controlling agent.

## Author contributions

AK conceived and designed the experiments. FF performed all the experiments. LV contributed in proteomics experiment. VMR contributed in growing the plants and quantitative gene expressions. GT provided resources for proteomics and supervised the sample preparation. AK and FF analyzed all the data, wrote and edited the manuscript by taking inputs from all the authors. All the authors read and approved the manuscript.

## Acknowledgements

We acknowledge BRIC-RGCB for intramural funding (RGCB/2023/00661 and RGCB/2025/01022) and University Grants Commission (UGC), India for funding the research fellowship (No. 231610103705) to Ms. Febina Fernandez. We acknowledge BRIC-RGCB Mass Spectrometry facility (funded by DBT-SAHAJ, Govt. of India) used for the service of LC-HRMS analysis and DBT-eLibrary Consortium (DeLCON) for providing access to e-resources.

## Data availability

Major data are available in the manuscript and supplementary files. Raw data will be available on request with valid reason.

## Supporting information

Fig. S1 Schematic diagram of ‘omics’ pipeline and sample’s integrity.

Fig. S2 Correlation in between top 500 metabolites and proteins fold change.

Fig. S3 KEGG pathway based metabolic pathway enrichment analysis with proteins.

Fig. S4 Proteomic and transcriptomics levels of anthocyanidin synthase (ANS) in control, 12 hpi and 24 hpi of *P. myriotylum* inoculated collar tissue samples of ginger.

Fig. S5 Metabolome alteration patterns in ginger collar in control, 12 hpi and 24 hpi upon *P. myriotylum* infection.

Fig. S6 KEGG pathway based metabolic chemical class enrichment analysis with metabolite features.

Fig. S7 Biosynthesis of the phenylpropanoid pathway associated major metabolites in ginger after *P. myriotylum* infection.

Fig. S8 Correlation of jasmonic acid with other defense metabolites.

Table S1. Oligonucleotide primers used for the RT-qPCR analysis and fungus specific primers for semiquantitative PCR

Table S2. Significantly altered proteins at 12 hpi

Table S3. Significantly altered proteins at 24 hpi

Table S4. Venn Analysis of proteome data

Table S5. Pathway Analysis with proteome data at 12 hpi and 24 hpi

Table S6. Top 20 enriched pathways at 12 hpi and 24 hpi

Table S7. Deconvoluted metabolite features at 12 hpi and 24 hpi

Table S8. Significantly altered metabolite features at 12 hpi and 24 hpi

Table S9. Annotated specialized metabolites associated with enriched secondary metabolic pathways at 12 hpi and 24 hpi

